# A novel consensus-based computational pipeline for rapid screening of antibody therapeutics for efficacy against SARS-CoV-2 variants of concern including omicron variant

**DOI:** 10.1101/2022.02.11.480177

**Authors:** Naveen Kumar, Rahul Kaushik, Kam Y. J. Zhang, Vladimir N. Uversky, Pratiksha Srivastava, Upasana Sahu, Richa Sood, Sandeep Bhatia

**Author notes:** Correspondence: E-mail address (Naveen Kumar). These authors contributed equally to this manuscript (Naveen Kumar, Rahul Kaushik).

## Abstract

Multiple severe acute respiratory syndrome coronavirus 2 (SARS-CoV-2) variants continue to evolve carrying flexible amino acid substitutions in the spike protein’s receptor binding domain (RBD). These substitutions modify the binding of the SARS-CoV-2 to human angiotensin-converting enzyme 2 (hACE2) receptor and have been implicated in altered host fitness, transmissibility and efficacy against antibody therapeutics and vaccines. Reliably predicting the binding strength of SARS-CoV-2 variants RBD to hACE2 receptor and neutralizing antibodies (NAbs) can help assessing their fitness, and rapid deployment of effective antibody therapeutics, respectively. Here, we introduced a two-step computational framework with three-fold validation that first identified dissociation constant as a reliable predictor of binding affinity in hetero-dimeric and –trimeric protein complexes. The second step implements dissociation constant as descriptor of the binding strengths of SARS-CoV-2 variants RBD to hACE2 and NAbs. Then, we examined several variants of concern (VOCs) such as Alpha, Beta, Gamma, Delta, and Omicron and demonstrated that these VOCs RBD bind to the hACE2 with enhanced affinity. Furthermore, the binding affinity of Omicron variant’s RBD was reduced with majority of the RBD-directed NAbs, which is highly consistent with the experimental neutralization data. By studying the atomic contacts between RBD and NAbs, we revealed the molecular footprints of four NAbs (GH-12, P2B-1A1, Asarnow_3D11, and C118) — that may likely neutralize the recently emerged omicron variant — facilitating enhanced binding affinity. Finally, our findings suggest a computational pathway that could aid researchers identify a range of current NAbs that may be effective against emerging SARS-CoV-2 variants.

## Introduction

The ongoing coronavirus disease 2019 (COVID-19) pandemic caused by severe acute respiratory syndrome-coronavirus-2 (SARS-CoV-2) continues to be a serious global public health issues that has claimed the lives of several million people. Over the course of COVID-19 pandemic, the SARS-CoV-2 Wuhan-Hu-1 or wild-type (WT) virus has acquired multiple mutations, resulting in emergence of multiple variants of concern (VOCs) such as alpha (B.1.1.7) in United Kingdom, beta (B.1.351) in South Africa, gamma (P.1) in Brazil, and delta (B.1.617.2) in India [1–4]. The World Health Organization (WHO) has defined these VOCs as having any of the following characteristics: increased transmissibility, increased virulence or change in clinical illness presentation, or decreased vaccination or treatment effectiveness. These VOCs have developed resistance to many neutralizing antibodies, including some clinical antibodies approved by the United States Food and Drug Administration (FDA) as therapeutics, rendering them ineffective for treatment of COVID-19 patients [5,6].

The newly emerged omicron (B.1.1.159) VOC in South Africa has the largest number of mutations — 30 amino acid substitutions, deletion of six residues, and insertion of three residues — in its spike protein relative to that of wild-type virus [7]. The receptor-binding motif (RBM) in spike protein that mediates binding of virus to the host cells via angiotensin-converting enzyme 2 (ACE2) receptor contains flexible amino acid substitutions (one mutation in alpha, three in beta and gamma, two in delta, and ten in omicron). These substitutions resulted in improved transmissibility by increased binding affinity to the ACE2 receptor [8,9] or reduced neutralization by post-immunization serum or neutralizing antibodies (NAbs), especially for omicron variant [10–12]. The majority of the existing NAbs have been found to be ineffective against the SARS-CoV-2 omicron variant [13]. Furthermore, with these VOCs circulating in humans, more variants are highly likely emerge in future with altered binding affinity to human ACE2 (hACE2) receptor [14] and may compromise the neutralizing ability of convalescent or post-immunization serum or neutralizing antibodies (NAbs).

In the event of the introduction of new SARS-CoV-2 VOCs, all NAbs should be tested experimentally against the new VOCs to discover the most effective NAbs that have retained their neutralizing ability, however this would be extremely labor-intensive, time-consuming and costly. In such scenario, computational methods that mimic the experimental binding affinity of SARS-CoV-2 spike protein to hACE2 receptor could be crucial in quickly understanding SARS-CoV-2 host-adaptability and guiding researchers toward employing a soluble hACE2-Fc decoy as a SARS-CoV-2 therapy. Furthermore, such methodologies could be used to screen a pool of current NAbs for efficacy against new VOCs and assist researchers in selecting NAbs for further neutralization testing in the lab.

To address this issue, we established a novel consensus-based computational pipeline — by benchmarking our findings with the experimental results — that reliably predicts binding affinity between the hACE2 receptor and the SARS-CoV-2 spike’s receptor-binding domain (RBD) (Figure 1). We also demonstrated that the binding affinity of SARS-CoV-2 spike’s RBD to a panel of NAbs estimated by our established computational pipeline was highly correlated with the experimental neutralization data, indicating the robustness and reliability of this approach for a rapid screening of current NAbs for efficacy against the new SARS-CoV-2 variant. Furthermore, we not only suggest four NAbs that may likely neutralize the recently emerged SARS-CoV-2 omicron variant, but we also examine the molecular footprints shared by these NAbs, which likely provided them with the enhanced binding affinity, by studying the atomic contacts between RBD and NAbs.

**Figure 1.**
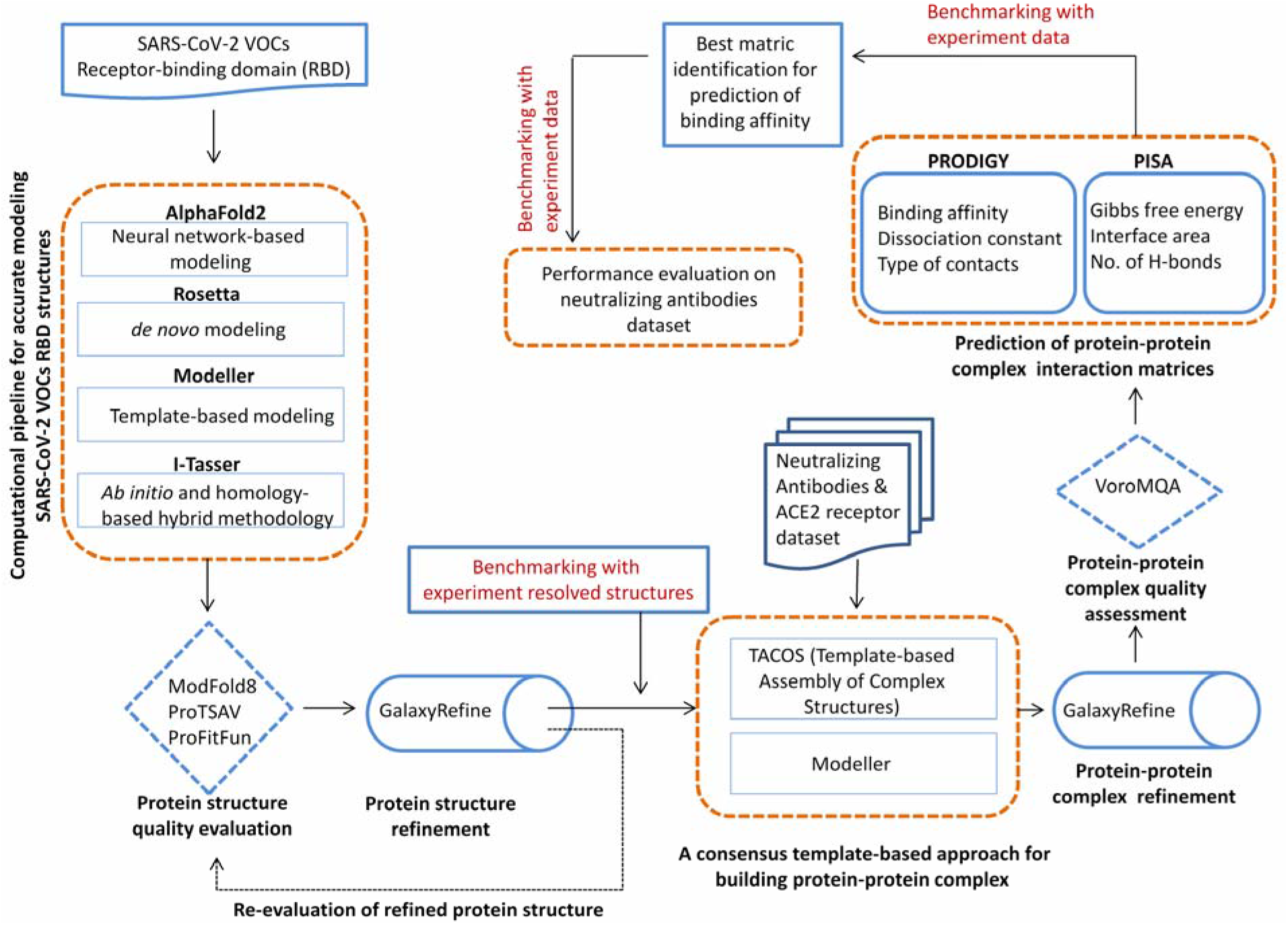
A schematic representation of the two-step computational framework with three-fold validation for rapid screening of antibody therapeutics against SARS-CoV-2 variants.

## Materials and Methods

### SARS-CoV-2 VOCs and NAbs dataset

In order to avoid the intrinsic bias in the data selection, we randomly selected the complete genomic sequences representing distinct SARS-CoV-2 VOCs, viz., alpha (GISAID accession no. EPI_ISL_679974), beta (GISAID accession no. EPI_ISL_2493065), gamma (GISAID accession no. EPI_ISL_978506), delta (GISAID accession no. EPI_ISL_1663516) and omicron (GISAID accession no. EPI_ISL_6854348) from the EpiCoV™ database maintained by the Global Initiative on Sharing All Influenza Data (GISAID) [15]. These multiple sequences were aligned by MAFFT [16] to extract the sequence fragment encoding for receptor-binding domain of spike protein (T333 – G526). These RBD protein sequences corresponding to SARS-CoV-2 VOCs were subsequently employed for computational modeling of their RBD structures.

The Coronavirus Antibody Database (CoV-AbDab, http://opig.stats.ox.ac.uk/webapps/covabdab/) sponsored by the University of Oxford maintains detailed information of all the neutralizing antibodies tested against the SARS-CoV-2 wild type virus [17]. We selected sixty-seven NAbs based on the criteria that (i) they target spike’s RBD, (ii) their experimental structures have been solved, and (iii) they do not have missing or non-standard residues especially at the RBD-NAb interfaces. Subsequently their protein structures were extracted from Protein Databank [18]. The details of selected NAbs including their source, germ line and their structures identity (PDB Id) is provided in Table S1.

### Computational pipeline for accurate modeling SARS-CoV-2 VOCs RBD structures

Recent advances in protein structure prediction have yielded several very reliable *de-novo*, knowledge-based and hybrid approaches that take advantage of the inherent familiarity with experimentally solved protein structures [19–24]. In this study, we used an exhaustive consensus computational technique to obtain highly trustworthy RBD protein structures of SARS-CoV-2 VOCs. The preliminary model structures for each of the SARS-CoV-2 VOCs RBD were predicted using four most promising state-of-the-art methods: I-Tasser (version 5.1) (*ab initio* and homology-based hybrid methodology) [24]; Modeller (version 10.1) (template-based modeling) [19]; standalone versions of Rosetta (*de novo* modeling) [23], and AlphaFold2 (available through Colab-Notebook) [20]. These algorithms, in particular, employ various approaches to protein structure prediction and have yielded 20 model structures for a single RBD sequence. To select the best model structure from each set of model structures, we used a consensus approach for protein structure quality evaluation by ModFold8 [25], ProTSAV [26], and ProFitFun [27,28]. On applying GalaxyRefine, a comprehensive protein structure refinement approach further improved the selected model structure for each RBD protein [29].

### Building RBD-NAbs complexes through a consensus template-based approach and their refinement

Finding adequate templates and determining the interacting protein partners for the target protein are, in general, the most significant bottlenecks for accurate modeling of protein-protein complexes. In our investigation, a template-based protein-protein docking was performed using TACOS (Template-based Assembly of Complex Structures) [30] as a standalone tool to predict 5 model structures of the RBD-NAb complex for each NAb. Additionally, we used template-based modeling to yield five more RBD-NAb complex structures for each NAb by taking corresponding experimentally solved RBD^WT^-NAb complex using Modeller (version 10.1) [19]. Eventaully, we ended up in generating ten RBD-NAb complex structures for each NAb.

To refine the RBD-NAb complexes (ten models for each NAb), we used standalone version of GalaxyRefineComplex [31] that resulted fifty models for each RBD-NAb complex. These protein complexes were assessed to determine the best refined and optimized RBD-NAb complex for each NAb by using protein-protein complex quality assessment through VoroMQA [32]. The best RBD-NAb protein-complex for each NAb was used to predict binding affinity in order to assess their ability to neutralize SARS-CoV-2 VOCs and also dissect the molecular interactions at the RBD-NAb complex interface.

### Prediction of binding affinity of SARS-CoV-2 spike’s RBD with hACE2 receptor and RBD-NAbs through different matrices and their benchmarking with experimental data

Diverse approaches for assessing protein-protein interactions in the protein complexes yield distinct metrics for quantifying their intermolecular interaction strength. Some of these metrics include predicted binding affinity (Kcal/mol), predicted dissociation constant (Kd), change in Gibbs free energy (ΔG), binding energy, interface area (Å), and various type of contacts or interactions such as hydrophobic-hydrophobic, polar-polar, polar-hydrophobic, H-bonds, and disulfide bonds. We employed the standalone versions of PRODIGY [33] and PISA [34] on the most reliable RBD-hACE2 and RBD-NAbs complexes to estimate these distinct matrices.

In addition, in order to discover the best metric that simulate with experimental binding affinity, a dataset comprised of hetero-dimeric (n = 400) and hetero-trimeric (n = 180) protein-protein complexes was retrieved from the PDBbind database, which provides detailed information on experimentally measured binding affinity for all biomolecular complexes deposited in the Protein Data Bank [35]. We employed Spearmen correlation statistical method to estimate the correlation between the experimental binding affinity data with the distinct matrices and demonstrated that predicted dissociation constant is the best metric that closely simulate with the experimental binding affinity. The summary of all the biomolecular complexes retrieved from the PDBbind database is provided in Table S2.

### Performance evaluation on a dataset of neutralizing antibodies (NAbs)

Following the demonstration that predicted dissociation constant is the most accurate metric for mimicking experimental binding affinity; it is unclear whether predicted binding affinity correlates with the experimental neutralization data. Therefore, we first predicted dissociation constant on the most reliable RBD-NAbs complex generated for each of sixty seven NAbs using PRODIGY [33], and subsequently evaluated the performance of this approach by comparing the experimental neutralization data derived from recently published studies [13,36]. We also dissect the molecular basis of enhanced binding affinity of four NAbs to SARS-CoV-2 omicron variant by analyzing the atomic contacts between RBD and NAbs.

### Statistical analysis

We employed Spearmen correlation statistical method to estimate the correlation between the experimental binding affinities data with the distinct matrices employed to assess the protein-protein interactions. The agreement between the predicted dissociation constant and experimental neutralization data was estimated by inter-rater reliability analysis using Kappa (κ) statistic [37]. Values of κ < 0.20 indicate poor agreement; 0.21 to 0.40, fair agreement; 0.41 to 0.60, moderate agreement; 0.61 to 0.80, substantial agreement and 0.81 to 1.00, perfect agreement. We employed MedCalc® statistical software version 20.026 to calculate weight Kappa statistic. The graphs were generated using GraphPad Prism 7.01 (GraphPad Software, San Diego, CA).

## Results and Discussion

Control of SARS-CoV-2 pandemic is only possible through generation of widespread population immunity acquired through vaccination or infection. Neutralizing antibodies (NAbs) in particular play crucial role in generating protective immunity against SARS-CoV-2. However, after almost two years into the pandemic, a considerable number of SARS-CoV-2 lineages are rising in frequency. These emerging lineages, particularly the variants of concern (VOCs), have acquired a few mutations in the SARS-CoV-2 spike protein’s receptor-binding domain (RBD), and these mutations often substantially sufficient to evade the protective immunity offered by the NAbs elicited either by infection or vaccination [38–40] (Figure 2a). In the case of the emergence new SARS-CoV-2 variant(s), such as recently emerged omicron VOC, the most promising therapeutic approach is finding the potent NAbs that have retained their neutralization activity against new SARS-CoV-2 variant(s). It would be highly time-consuming, expensive, and labor-intensive to experimentally test all of the existing NAbs against the new SARS-CoV-2 variant(s). To address this issue, we have established a novel consensus-based computational pipeline that reliably predict binding affinity in hetero-dimeric and hetero-trimeric protein complexes, which will be crucial not only in gaining a rapid understanding of SARS-CoV-2 host-adaptability, but also in guiding researchers in quickly selecting potent NAbs against the new SARS-CoV-2 variant(s) from a set of existing NAbs.

**Figure 2.**
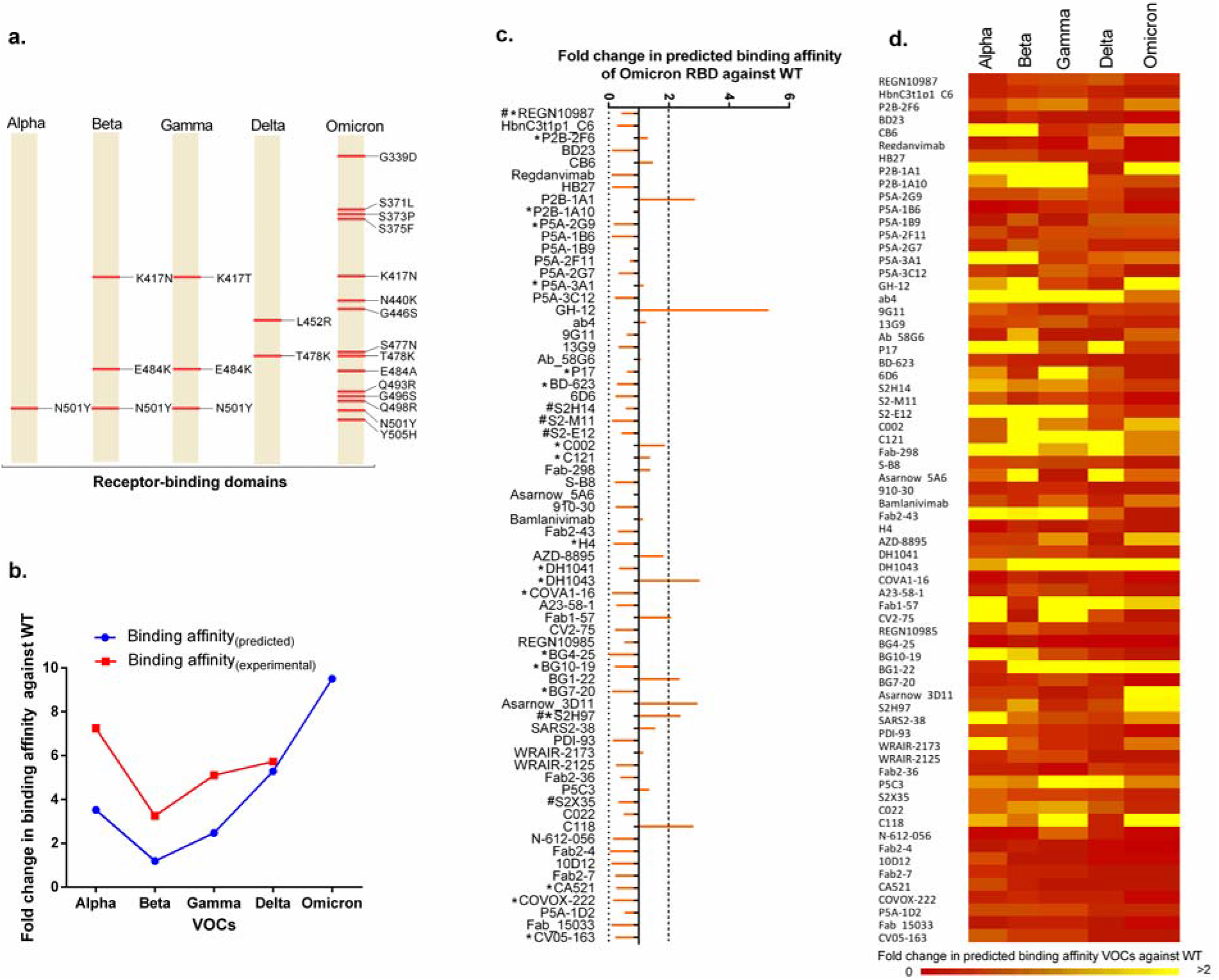
(a) A representation of the amino acid substitutions in the receptor-binding domains (RBDs) of SARS-CoV-2 VOCs, (b) Prediction of binding affinity of SARS-CoV-2 RBD with hACE2 receptor and its comparison with the experimental binding affinity, (c) the fold change in the predicted binding affinity of SARS-CoV-2 omicron VOC with a panel of neutralizing antibodies in comparison to wild type virus. # and * represent the neutralizing antibodies tested by Cameroni et al., 2021 and Cao et al., 2021, respectively, for their neutralization potential against SARS-CoV-2 omicron VOC, (d) A heat map showing the fold change in predicted binding affinity of SARS-CoV-2 VOCs with a panel of neutralizing antibodies in comparison to wild type virus.

Importantly, majority of neutralizing activity of polyclonal antibody response generated on account of SARS-CoC-2 infection is due to antibodies that target RBD, however, antibodies that target spike’s N-terminal domain (NTD) also contribute to neutralization [41–43]. These NAbs targeting the RBD or NTD of the spike protein offer protections by inhibiting the attachment of SARS-CoV-2 spike protein to ACE2 receptor on the target cells. Experimental studies have shown that the most potent NAbs frequently targets receptor-binding motif, located within the RBD [42,44]. Therefore, we first generated RBD model structures for all the SARS-CoV-2 VOCs and subsequently assess their quality by comparing with their experimentally solved structures (alpha: 7EKF, beta: 7EKG, gamma: 7EKC, delta: 7V8A, and omicron: 7T9L). The results showed that the root mean squared deviation of the modeled RBD structures to that of experimental counterpart is very low (alpha: 0.36 Å, beta: 0.55 Å, gamma: 0.27 Å, delta: 0.43 Å, and omicron: 0.85 Å), implying that the computational pipeline used in our study is suitable for prediction of RBD structures for emerging and new SARS-CoV-2 variants, whose experimental structure has not been determined. Hence, this integrated computational strategy for protein structure prediction and refinement ensured highly reliable protein modelled structures suitable for further downstream applications.

Different computational methods may produce different metrics for prediction of binding affinity in protein-protein complexes, and the resulting mispredictions might influence the reliability of subsequent analyses and interpretations. Thus, to find the best metric that closely matches with experimental binding affinity, we benchmarked predicted dissociation constant (Kd), and change in Gibbs free energy (ΔG) against experimental binding affinity on a dataset of hetero-dimeric (n = 400) and hetero-trimeric (n = 180) protein-protein complexes retrieved from the PDBbind database. The results revealed that the predicted dissociation constant is significantly highly correlated with the experimental binding affinity data (r = 0.858, P < 0.0001 for hetero-dimeric; r = 0.706, P < 0.0001 for hetero-trimeric protein complexes), implying that predicted dissociation constant is the most reliable metric to assess the binding affinity in both hetero-dimeric and hetero-trimeric protein complexes. Subsequently, we modeled RBD-hACE2 protein complex models for all the SARS-CoV-2 VOCs and compared their predicted binding affinity (via dissociation constant) with their experimental counterparts. We demonstrated that the SARS-CoV-2 VOCs RBD has increased binding affinity (≈3.5, ≈1.2, ≈2.5, and ≈5.3– fold higher for alpha, beta, gamma, and delta, respectively) to hACE2 receptor, which is consistent with the experimental binding affinity estimated by surface plasmon resonance [9] (Figure 2b). Our findings also revealed that the omicron RBD has a higher predicted binding affinity to the hACE2 receptor (≈9.5–fold greater than wild type or Wuhan-Hu-1 strain), which is in concordance with the experimental binding affinity (≈3 – fold higher) estimated in a recent study [36]. Furthermore, we noted the differences in the experimental binding affinity estimation between studies [9,36] that could be related to the use of various wild type or VOCs viruses in the diverse experimental lab setups. However, the findings of both of these studies, including our study, consistently demonstrated that the SARS-CoV-2 VOCs including omicron variant have a higher binding affinity for the hACE2 receptor. Estimating the binding affinity of SARS-CoV-2 spike protein to the ACE2 receptor has direct implications not just for identifying the host range of SARS-CoV-2 [45], but also for adaptation and infection dynamics in the host [40].

It’s worth noting that the potent NAbs have a high affinity for RBD, implying a positive correlation between neutralization ability and binding affinity [46]. Consequently, we implemented the computational pipeline developed in our study for the prediction of binding affinity of SARS-CoV-2 VOCs RBD with a panel of NAbs retrieved from the Coronavirus Antibody Database (CoV-AbDab, http://opig.stats.ox.ac.uk/webapps/covabdab/) [17] and subsequently, to validate our approach, we compared the predicted binding affinity results with the experimental neutralization data. A total of 67 NAbs were chosen based on the criteria that they target spike’s RBD and their experimental structures have been solved. To reduce the false positive results, we adjusted the cut-off to at least 2-fold increment in binding affinity against the wild type virus for NAbs to be designated as effective neutralization of Omicron VOC. Based on this criteria, the majority of NAbs (about 90%, 60 out of 67) were not effective against SARS-CoV-2 Omicron VOC, which is consistent with recent studies showing that the majority of NAbs are ineffective against Omicron VOC [13,36] (Figure 2c, Table S3). Furthermore, none of the NAbs can be classified as broadly-acting against all the SARS-CoV-2 VOCs (alpha, beta, gamma, delta, and omicron) (Figure 2d). On comparing with the experimental results of both studies, our findings are in perfect agreement with Cameroni et al., 2021 (weighted kappa κ = 1; 6/6), and good agreement with Cao et al., 2021 (weighted kappa κ = 0.643, 95% CI= 0.0055778 to 1.00000; 19/20). Furthermore, Regeneron’s REGN-COV2 antibody cocktail (REGN10933 and REGN10987; which has been granted for emergency use authorization for antibody treatment for COVID-19) is ineffective against Omicron VOC, which is in line with experimental neutralization assays (Figure 2c) [13,36], suggesting that this computational pipeline is also suitable for screening the antibody cocktail for their efficacy against SARS-CoV-2 variants. Our results remained consistent when we estimated the binding affinities of REGN10933 and REGN10987 to RBD individually (Table S3). Finally, in our investigation, four NAbs, GH-12, P2B-1A1, Asarnow_3D11, and C118, showed increased predicted binding affinity to omicron VOC RBD compared to wild-type SARS-CoV-2.

The NAbs can be broadly classified into two categories, those that inhibit the interaction of spike protein with the ACE2 receptor by binding to epitopes that share footprints with receptor-binding motif (RBM), and those that bind to non-RBM epitopes and thus do not block the interaction of spike protein with the ACE2. The most potent NAbs frequently target RBM, which directly interferes with the binding of ACE2. However, neutralizing potential of these NAbs gets abrogated on account of accumulation of multiple mutations during the SARS-CoV-2 evolution [13,36,47]. Therefore, we investigated the binding sites of 4 NAbs (GH-12, P2B-1A1, Asarnow_3D11, and C118) on omicron spike protein and compared the molecular interactions with S2H97 (NAb effective against omicron) and REGN-COV2 antibody cocktail (REGN10933 and REGN10987). Of four NAbs, three (GH-12, Asarnow_3D11, and C118) bind to non-RBM epitopes similar to that of S2H97, while P2B-1A1 binding epitopes share footprints with that of REGN10987 by binding to the RBM (Figure 3a-h). Both the heavy and light chains of S2H97 (13 polar contacts and 6 salt bridges), GH-12 (12 polar contacts and 2 salt bridges), Asarnow_3D11 (11 polar contacts and 1 salt bridge), and C118 (18 polar contacts and 4 salt bridges) bind to a highly conserved cryptic antigenic site on RBD, designated previously as site V [47]. Although the binding epitope footprints of P2B-1A1 and REGN-COV2 antibody cocktail are similar, P2B-1A1 generated more polar interactions than the wild type virus (Figure 3a, Table S4). Furthermore, REGN-COV2 antibody cocktail lost the neutralization potential against omicron because it made entirely distinct and fewer polar interactions with omicron RBM (ASN417, LYS444, VAL445, TYR449, ASN450, ARG493, TYR501) than wild type virus (ASN440, VAL445, ASN448, TYR449, TYR453, GLU484, GLY485, PHE486, CYS488, TYR489, GLN498), and also lost one salt bridge due to LYS417ASN in omicron RBD (Table S4). Largely, our findings revealed that four NAbs, GH-12, P2B-1A1, Asarnow_3D11, and C118, have an enhanced binding affinity to omicron RBD than wild type SARS-CoV-2, indicating that further experimental research is needed to determine their neutralization potential.

**Figure 3.**
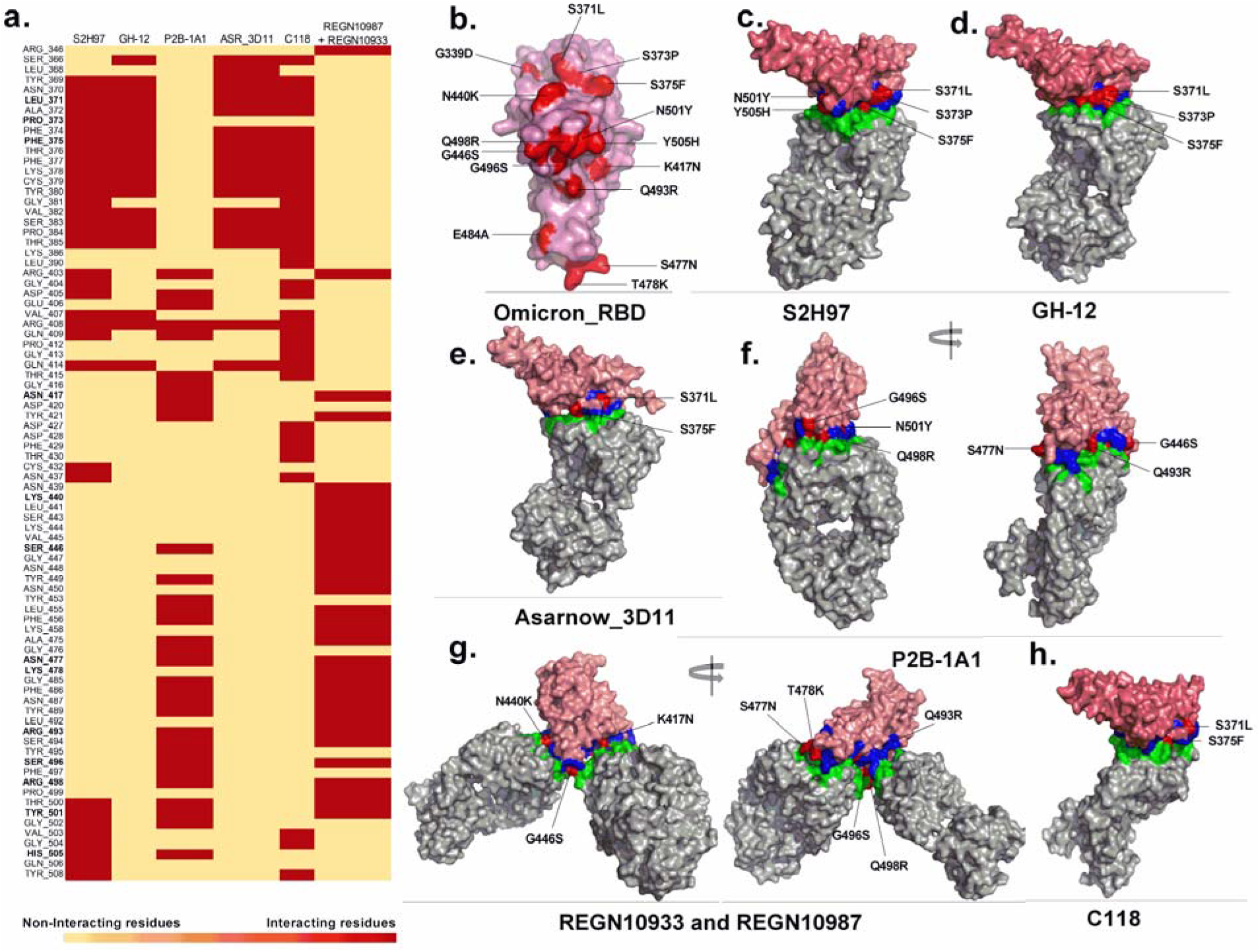
Structural insights into differential binding affinities of selected neutralizing antibodies to SARS-CoV-2 omicron VOC. (a) A heat map depiction of molecular interactions at the RBD-NAbs interface for selected NAbs (S2H97, GH-12, PB2-1A1, Asarnow_3D11, C118, REGN10987, and REGN10933), (b) the receptor-binding domain (RBD) of SARS-CoV-2 omicron VOC showing amino acids substitutions (indicated in red color) in comparison to wild type SARS-CoV-2, (c-h) SARS-CoV-2 omicron VOC RBD bound to S2H97 NAbs (c, PDB Id: 7M7W); GH-12 (d, PDB Id: 7D6I); Asarnow_3D11 (e, PDB Id: 7KQB); P2B-1A1 (f, PDB Id: 7CZP); REGN10933 and REGN10987 (g, PDB Id: 6XDG); and C118 (h, PDB Id: 7RKV). The interacting residues at the interface of RBD-NAbs are colored blue and green, respectively. The residues changed in Omicron RBD with respect to wild type RBD at the RBD-NAbs interface are red-colored.

## Conclusion

We introduced a three-fold validated two-step computational methodology for predicting binding affinity in hetero-dimeric and -trimeric protein complexes. This method is appropriate for quickly determining SARS-CoV-2 host-adaptability, screening existing NAbs for efficacy against current and new SARS-CoV-2 variants, and assisting researchers in selecting NAbs for further experimental neutralization testing. The NAbs could be tailored by targeting the conserved epitopes on spike’s RBD, particularly outside the RBM and utilizing a cocktail of such tailored NAbs as therapeutic drugs would be critical in combating current and emerging SARS-CoV-2 variants, as well as limiting the generation of SARS-CoV-2 escape mutants.

## Acknowledgments

The authors acknowledge the Advanced Center for Computing and Communication (ACCC) of the Institute of Physical and Chemical Research (RIKEN) for computing resources on the Hokusai BigWaterfall supercomputer. Finally, all of the authors thank and express gratitude to their respective Institutes.

## Data Availability Statement

All the datasets related to this study have been provided as supplementary information and can be accessed online at https://academic.oup.com/bib.

